# Management of Mendelian Traits in Breeding Programs by Gene Editing: A Simulation Study

**DOI:** 10.1101/116459

**Authors:** John B. Cole

**Author notes:** Email addresses: JBC.

## Abstract

**Background:** Genotypes based on high-density single nucleotide polymorphisms have recently been used to identify a number of novel recessive mutations that adversely affect fertility in dairy cattle as well as to track conditions such as polledness. The use of sequential mate allocation strategies that account for increases in genomic inbreeding and the economic impact of affected matings may result in faster allele frequency changes than strategies that do not consider inbreeding and monetary losses. However, the effect of gene editing on selection programs also should be considered because gene editing has the potential to dramatically change allele frequencies in livestock populations.

**Methods:** A simulation program developed to evaluate dairy cattle breeding schemes was extended to include the use of clustered regularly interspaced short palindromic repeat (CRISPR), transcription activator-like effector nuclease (TALEN), and zinc finger nuclease (ZFN) technologies for gene editing. A hypothetical technology with a perfect success rate was used to establish an upper limit on attainable progress, and a scenario with no editing served as a baseline for comparison.

**Results:** The technologies differed in the rate of success of gene editing as well as the success rate of embryo transfer based on literature estimates. The number of edited alleles was assumed to have no effect on success rate. The two scenarios evaluated considered only the horned locus or 12 recessive alleles that currently are segregating in the U.S. Holstein population. The top 1, 5, or 10% of bulls were edited each generation, and either no cows or the top 1% of cows were edited. Inefficient editing technologies produced less cumulative genetic gain and lower inbreeding than efficient ones. Gene editing was very effective at reducing the frequency of the horned haplotype (increasing the frequency of polled animals in the population), and allele frequencies of the 12 recessives segregating in the U.S. Holstein population decreased faster with editing than without.

**Conclusions:** Gene editing can be an effective tool for reducing the rate of harmful alleles in a dairy cattle population even if only a small proportion of elite animals are modified.

## Background

The widespread adoption and corresponding reduction in the cost of high-density single nucleotide polymorphism (SNP) genotyping has enabled the detection of many new recessives that have deleterious effects on fertility in several breeds of dairy cattle [1,2,3], and whole genome sequencing allows detecting additional fertility defects [4]. Many of these new recessives were not previously detected by test matings because they cause embryonic losses in early gestation that could not be distinguished from failed breedings. Annual losses to U.S. dairy farmers from decreased fertility and increased perinatal mortality due to known recessive defects are estimated to be at least $10 million (€9,370,754) [3]. Mate allocation tools do not always consider carrier status when bull and cow pairs are assigned, and few make use of DNA marker or haplotype information. Avoiding carrier-to-carrier matings is easy when only a few recessives are segregating in a population but is considerably more difficult when many defects are segregating.

Cole [5] recently extended a simple method for controlling the rate of increase in genomic inbreeding proposed by Pryce et al. [6] to account for economic losses attributable to recessive defects. In the original method, parent averages (PAs) for matings that produced inbred offspring were penalized, and the bull that produced the highest PA after the inbreeding adjustment was selected in a sequential manner. The number of matings permitted for each bull was constrained to prevent one bull with high genetic merit from being mated to all cows. Cole [5] modified this approach to include an additional term that penalized carrier-to-carrier matings that may produce affected embryos and showed that the additional penalty decreased minor allele frequency (MAF) faster than other methods. However, many generations of selection were still needed to eliminate recessives from the population, and some defects remained in the population at low frequency.

A number of tools are now available for editing eukaryotic genomes, including clustered regularly interspaced short palindromic repeats (CRISPR), transcription activator-like effector nucleases (TALEN), and zinc finger nucleases (ZFN) [7,8]. Treating simple recessive disorders by using gene editing is of great interest (e.g., [9]), and CRISPR has been used to generate pigs that are resistant to porcine reproductive and respiratory syndrome [10]. Gene editing also has been used to produce desirable phenotypes (e.g., polled cattle [11]). A recent series of simulation studies showed that gene editing also has the potential to improve rates of genetic gain for quantitative traits [12,13]. Gene editing may be an effective means of reducing the frequency of genetic disorders in livestock populations or eliminating those disorders altogether.

The objective of this research was to determine rates of allele frequency change and quantify differences in cumulative genetic gain through simulation for several genome editing technologies while considering varying numbers of recessives and different proportions of bulls and cows to be edited.

## Methods

### Simulation

The simulation software of Cole [14] was modified to include four different gene editing technologies and used to examine several scenarios for the use of gene editing in a dairy cattle population. With the exception of the gene editing methodology, the simulation procedures were identical to those described in detail by Cole [5]. Thirteen software parameters were used in the simulations (Table 1).

**Table 1.** Simulation parameters.

| Software parameter | Definition | Value |
| --- | --- | --- |
| base_bulls | Number of bulls in the base population | 350 |
| base_cows | Number of cows in the base population | 35,000 |
| service_bulls | Number of bulls in the sire portfolio used by each herd | 50 |
| base_herds | Number of pseudo-herds used in the simulation | 200 |
| max_bulls | Maximum number of bulls available for use as service sires in each generation | 500 |
| max_cows | Maximum number of cows in the population in each generation | 100,000 |
| generations | Number of generations simulated | 20 |
| max_matings | Maximum number of matings each service sire is permitted each year | 5000 |
| debug | Show or hide debugging messages | True |
| history_freq | Frequency with which history files are saved to disk | End |
| rng_seed<sup>2</sup> | Value used to seed the random number generator | Time + PID |
| edit_prop | Proportions of bulls and cows edited in different scenarios | 0%, 1%, 10% (bulls);<br>0%, 1% (cows) |
| edit_type<sup>3</sup> | Technologies used for gene editing | C, P, T, Z |

*Time* system clock time when the simulation is submitted, *PID* process identification reported by the operating system, *C* clustered regularly interspaced short palindromic repeats, *T* transcription activator-like effector nuclease, *P* hypothetical technology with perfect success rate, *Z* zinc finger nuclease

### Mate allocation

The modified Pryce scheme accounting for recessive alleles described by Cole [5] was used to allocate bulls to cows in all scenarios. The selection criterion was the 2014 revision of the lifetime net merit (NM$) genetic-economic index used in the United States [15]. For each herd, 20% of the bulls were randomly selected from a list of live bulls, and the top 50 bulls from that group were selected for use as herd sires based on true breeding value (TBV). This produced different sire portfolios for each herd and is similar to the approach of Pryce et al. [6].

As in Cole [5], a matrix of PAs (**B***’*) was constructed with rows corresponding to bulls and columns corresponding to cows as

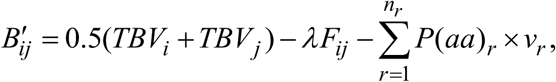

where *B’_ij_* is the PA for offspring of bull *i* and cow *j*, *TBV_i_* is the TBV NM$ for bull *i, TBV_j_* is the TBV NM$ for cow *j*, *λ* is the inbreeding depression in dollars associated with a 1% increase in inbreeding, *F_ij_* is the pedigree coefficient of inbreeding of the calf resulting from mating bull *i* to cow *j, n_r_* is the number of recessive alleles in a scenario, *P*(*aa*)_*r*_ is the probability of producing an affected calf for recessive locus *r*, and *v_r_* is the economic value of locus *r*. The regression coefficient of NM$ on inbreeding (*λ*) was computed as the weighted average of the December 2014 effects of inbreeding on the traits in the index as done by Cole [5]; the weights correspond to those assigned to each trait in the NM$ index and resulted in a *λ* of $25. The *P*(*aa*) equals 0.25 for a mating of two carriers, 0.5 for a mating of an affected animal with a carrier, or 1 for a mating of two affected animals. Thirteen recessive loci were used in the simulations (Table 2).

**Table 2.** Properties of the recessive loci included in each simulated scenario.

| Scenario | N<sup>a</sup> | Frequency | Value (\$) <sup>b</sup> | Name | Lethal |
| --- | --- | --- | --- | --- | --- |
| All recessive loci | 12 | 0.0276 | 150 | Brachyspina | Yes |
|  |  | 0.0192 | 40 | HH1 | Yes |
|  |  | 0.0166 | 40 | HH2 | Yes |
|  |  | 0.0295 | 40 | HH3 | Yes |
|  |  | 0.0037 | 40 | HH4 | Yes |
|  |  | 0.0222 | 40 | HH5 | Yes |
|  |  | 0.0025 | 150 | BLAD | Yes |
|  |  | 0.0137 | 70 | CVM | Yes |
|  |  | 0.0001 | 40 | DUMPS | Yes |
|  |  | 0.0007 | 150 | Mulefoot | Yes |
|  |  | 0.9929 | 40 | Horned | No |
|  |  | 0.0542 | -20 | Red coat color | No |
| Horned locus | 1 | 0.9929 | 40 | Horned | No |

*HH1, HH2, HH3, HH4, HH5* Holstein fertility haplotypes 1,2,3,4,5, respectively, *BLAD* bovine leukocyte adhesion deficiency, *CVM* complex vertebral malformation, *DUMPS* deficiency of uridine monophosphate synthase
<sup>a</sup>Number of recessive loci in the scenario.
<sup>b</sup>Positive values are undesirable and negative values are desirable.

After **B***’* was constructed, a matrix of matings (**M**) was used to allocate bulls to cows. An element (*M_ij_*) was set to 1 if the corresponding *B’_ij_* value was the greatest value in column *j* (that bull produces the largest PA of any bull available for mating to that cow); all the other elements of that column were set to 0. If the sum of the elements of row *i* was less than the maximum number of permitted matings for that bull, then the mating was allocated. Otherwise, the bull with the next-highest *B’_ij_* value in the column was selected. This procedure was repeated until each column had only one element equal to 1.

### Gene editing

In the simulation model, gene editing occurred when an embryo was created. The following six steps were used and repeated for each locus to be edited:

Step 1: Sort candidates on TBV in descending order.
Step 2: Select animals to be edited based on the user-specified proportion.
Step 3: Edit *Aa* and *aa* genotypes to *AA* genotypes (all edited animals are assumed to be homozygous).
Step 4: Draw a uniform random variate and compare with the editing failure rate of the method to determine if the editing procedure was successful. This check was made to determine if the recessive (*a*) alleles were successfully changed to dominant (*A*) alleles in the embryo.
Step 5: Draw a uniform random variate and compare with the embryonic death rate of the method to determine if the embryo transfer (ET) procedure was successful. This check was made to determine if the edited embryo survived the ET procedure and resulted in a live calf.
Step 6: Update the animal record.

The overall success rate was the product of the editing success and embryonic death rates (Steps 4 and 5). Figure 1 shows a flowchart describing the process in detail. The editing failure rate can be set to 0 to represent a scenario in which only embryos that were successfully edited are transferred to recipients. A scenario in which many embryos are produced so that survival of some is guaranteed can be simulated by setting the embryonic death rate to 0.

**Figure 1.**
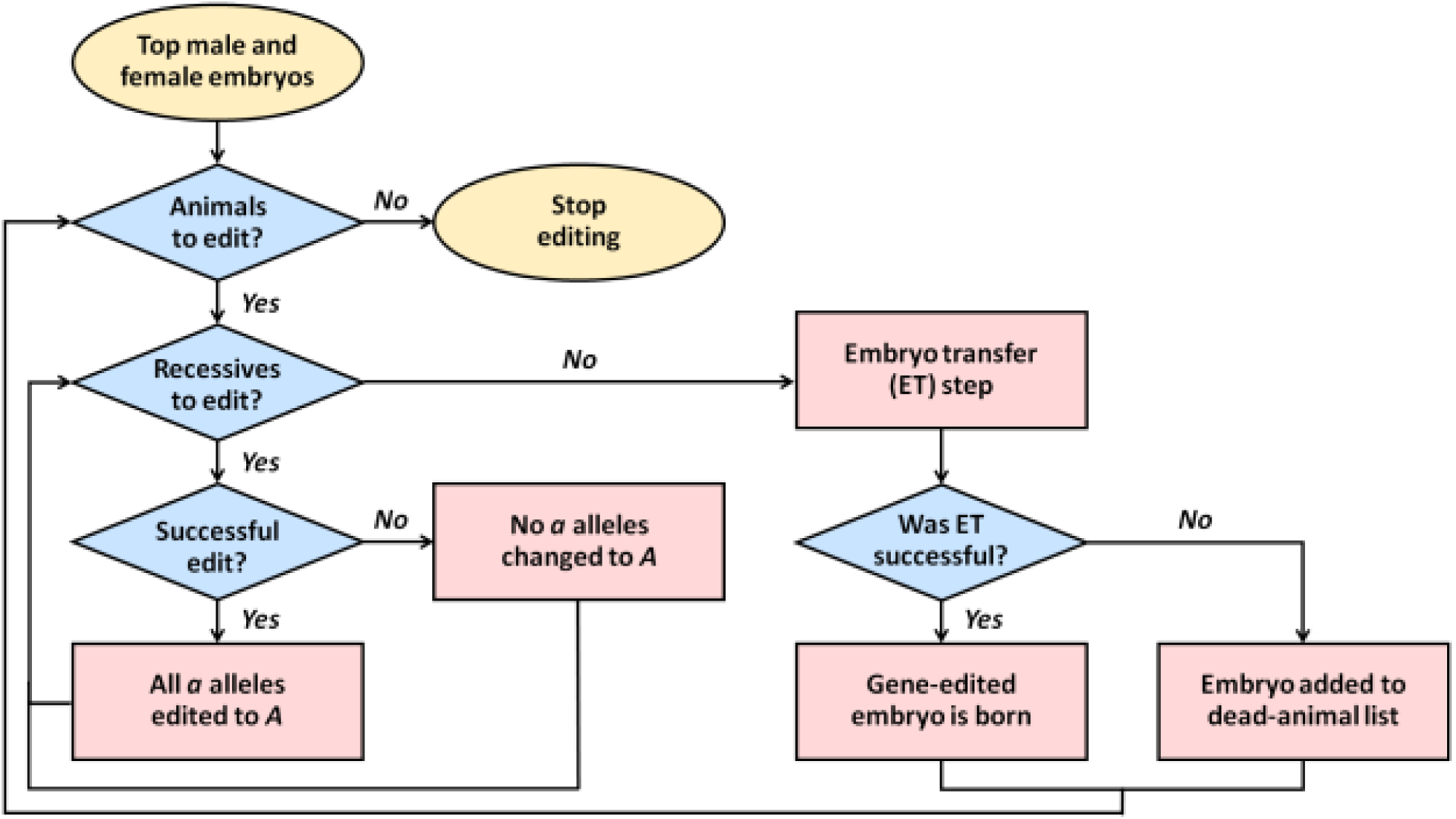
Flowchart for gene editing and embryo transfer in the simulation.

Three laboratory approaches to gene editing (CRISPR, TALEN, and ZFN) were supported as well as a fourth method that assumes that editing always is successful. The CRISPR, TALEN, and ZFN methods differed in their editing success and embryonic death rates [7,8] (Table 3). Bulls and cows could be edited at different rates (e.g., 10% of bulls and 1% of cows). Any combination of loci could be edited, and the number of edited loci was not restricted. A scenario in which no genes were edited, which reflects current practice, was used as the baseline against which the various editing scenarios were compared. A schematic of the simulated scenarios is in Figure 2.

**Table 3.**
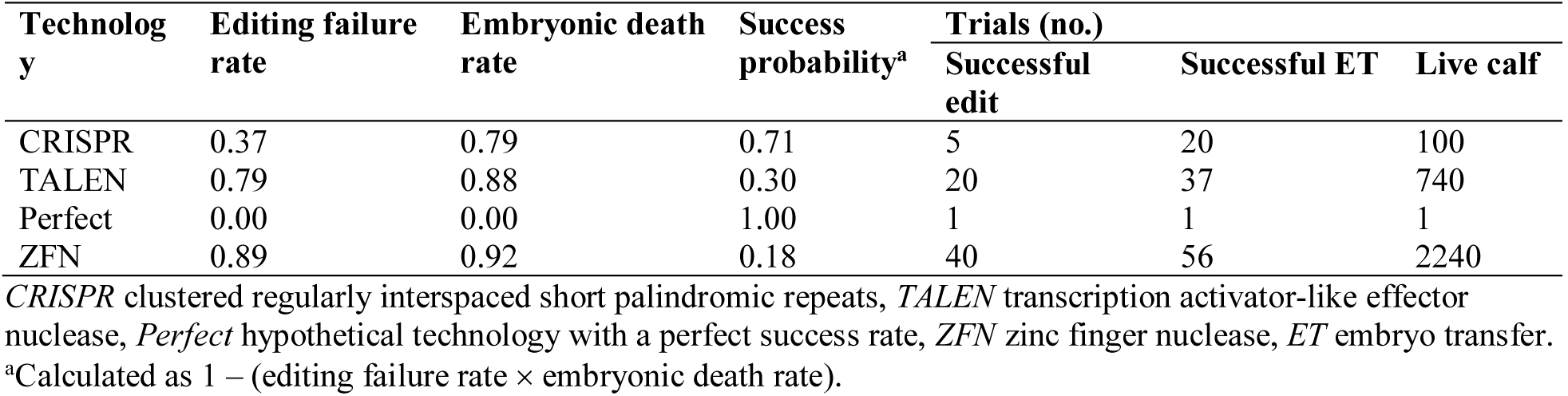
Gene editing failure and embryonic death rates and trials needed for a live calf.

| Technology | Editing failure rate | Embryonic death rate | Success probability<sup>a</sup> | Trials (no.) |  |  |
| --- | --- | --- | --- | --- | --- | --- |
|  |  |  |  | Successful edit | Successful ET | Live calf |
| CRISPR | 0.37 | 0.79 | 0.71 | 5 | 20 | 100 |
| TALEN | 0.79 | 0.88 | 0.30 | 20 | 37 | 740 |
| Perfect | 0.00 | 0.00 | 1.00 | 1 | 1 | 1 |
| ZFN | 0.89 | 0.92 | 0.18 | 40 | 56 | 2240 |

<sup>a</sup>Calculated as $1 - (\text{editing failure rate} \times \text{embryonic death rate})$ .

**Figure 2.**
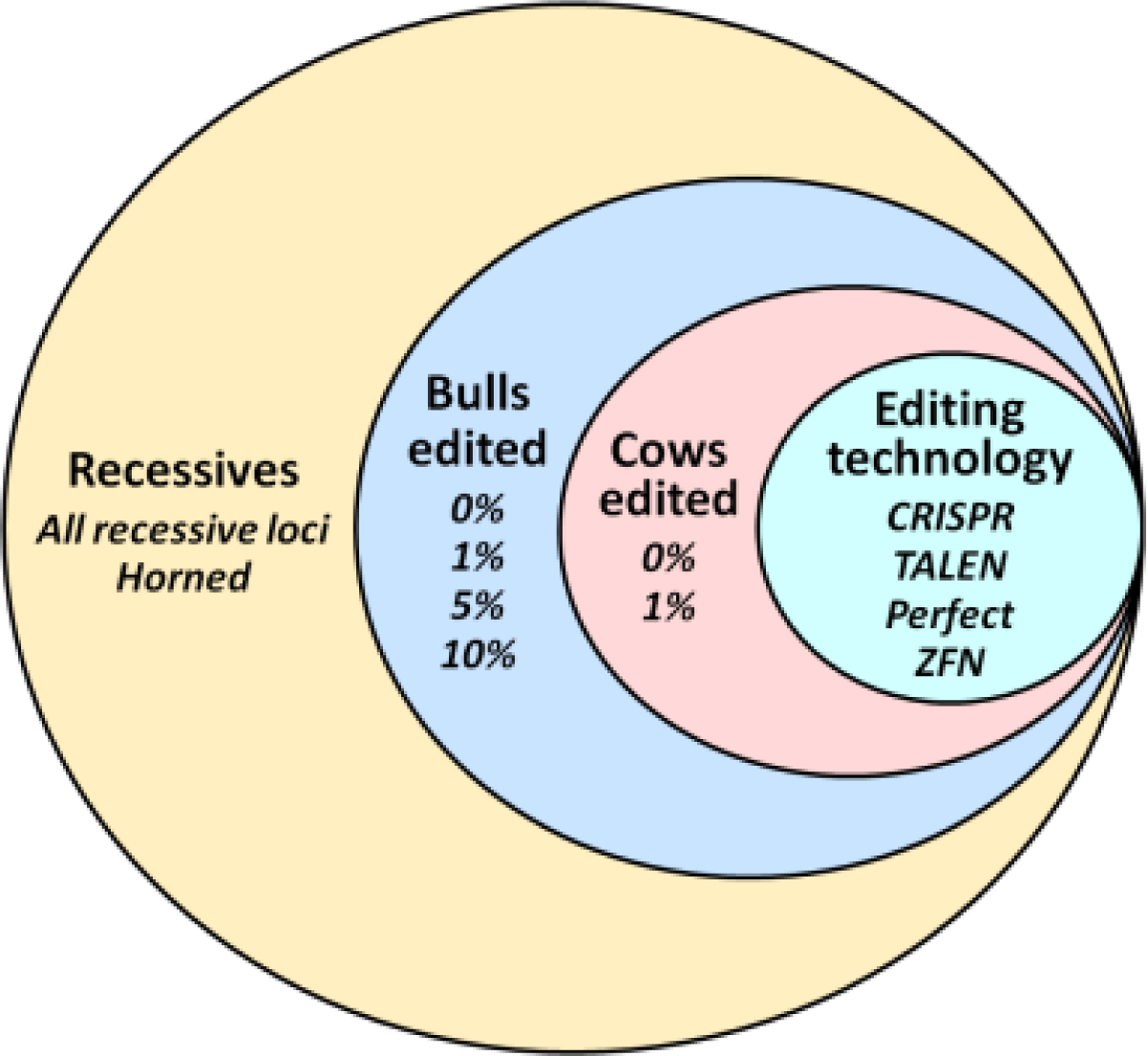
Schematic of simulated scenarios. Terms are nested within one another from left to right (e.g., the proportion of bulls edited is nested within the recessive scenario). *CRISPR* clustered regularly interspaced short palindromic repeats, *TALEN* transcription activator-like effector nuclease, *Perfect* = hypothetical technology with a perfect success rate, *ZFN* zinc finger nuclease

### Analysis

#### Trials required

The number of trials required to produce a live, gene-edited calf was determined for each of the four editing technologies (Table 3) by computing the number of draws needed from a geometric distribution to have a 99% probability of obtaining a success using the editing failure and embryonic death rates as the probability of success. The total number of trials was the product of the number of trials required for a successful edit and the number of trials needed for a successful ET. Producing a calf of the desired sex was assumed to be possible through the use of sexed semen, selection among the embryos in a flush, or other assisted reproductive technology.

#### Expected allele frequencies

The results for each scenario were averaged over 10 replicates. Observed changes in allele frequency were compared against expectations, and expected allele frequencies in each generation for lethal defects were calculated as in [16]:

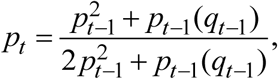

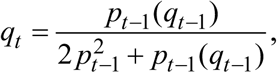

Where *p_t_* is the frequency of the major allele at time *t, q_t_* is the MAF at time *t*, and *t* ranges from 1 to 20 years. The MAF at time 0 was used in each scenario for each recessive locus (Table 2), and the major allele frequency was calculated as 1 – MAF. Expected frequencies for non-lethal alleles were calculated using Hardy–Weinberg proportions [17]:

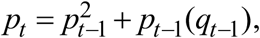

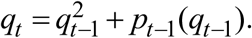

#### Rate of allele frequency change

For each recessive locus in each scenario, observed allele frequencies were regressed on birth year using the Python module Statsmodels version 0.6.1 ([18,19]) using the model:

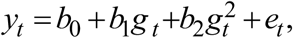

where *y_t_* is the frequency of a recessive locus at time *t*, *b*_0_ is the intercept, *b*_1_ is the regression coefficient associated with the linear effect of time, *g_t_* is the generation number at time *t*, *b*_2_ is the regression coefficient associated with the quadratic effect of time, *g_t_^2^* is the square of the generation number at time *t,* and *e_t_* is the random residual error.

#### Visualization

Plots of actual versus expected allele frequencies and the change in carrier proportions over time were constructed using matplotlib version 1.5.1 [20,21]. Changes in observed allele frequencies over time were plotted using Seaborn version 0.5.1 [22].

## Results and discussion

### Scenarios

Two scenarios are discussed: 1% of bulls and 0% of cows edited, and 10% of bulls and 1% of cows edited. These scenarios represented the two extremes (least versus most editing), and the results from the other scenarios were intermediate to these results. When selection is based on TBV and not carrier status, more efficient editing procedures generally produce greater responses. Recent research has shown that biopsies of bovine embryos, such as might be used for genotyping, do not affect pregnancy rate [23]; therefore, success rates might be improved through more rigorous ET protocols for edited embryos even when editing technologies differ. Although the cost of producing gene-edited animals decreases as the technology becomes more efficient, this study did not examine those differences because no data on actual costs of production were publicly available.

### Trials required for successful procedures

The numbers of trials required to ensure a 99% chance of successfully editing embryos (Step 4) and of getting a live calf on the ground following ET (Step 5) are in Table 3. Of the existing technologies, CRISPR was the most efficient by a factor of ~7, requiring only 100 trials to produce a live calf. ZFN was only a quarter as efficient, requiring 2240 trials to produce a live calf. Although determining the actual cost of producing a gene-edited calf is difficult, $10,000 per animal seems reasonable [24]. Production costs would then range from $1 million (CRISPR) to $22.4 million (ZFN). Such high costs would almost certainly make gene editing commercially non-viable in the scenarios considered in this study, but an increase in ET success rate to 50% [25] would reduce costs by a factor of 3 to 8. The cost of producing a live calf using CRISPR would then be only $350,000, which could easily be recovered from semen sales. If sexed semen is not used to ensure that a calf of the desired gender is born, then the totals should be doubled. For this analysis, only a single recessive was assumed to be edited, and more trials will be required if many loci are edited in the same embryo.

An additional assumption was that only one embryo was edited per mating (only a single trial was carried out). However, in practice, many embryos would be edited and transferred to ensure the live birth of a calf of the desired sex. As discussed, such cases may be simulated by setting the editing failure rate and the embryonic death rate to 0. Therefore, the results of this study are underestimates of allele frequency changes that might be observed in commercial production.

### Allele frequency changes

#### Recessive loci

Only results for HH3 from the simulations that included all 12 Holstein recessives and for horned are discussed. The other loci in the 12-recessive simulation had results similar to those for HH3 (for lethals) and horned (the polled locus). These trends are broadly similar to the results of Segelke et al. [26], who showed that MAFs decrease much faster when dams are selected using an index based on six loci (HH1, HH2, HH3, HH4, HH5, and polled) rather than on breeding values for fertility.

### HH3

The causal variant associated with HH3 is a non-synonymous mutation in the *SMC2* (*structural maintenance of chromosomes 2*) gene at 95,410,507 bp on bovine chromosome (BTA for *Bos taurus*) BTA8 [27] and currently has the highest allele frequency of any known deleterious recessive in U.S. Holsteins (0.0295). As technology efficiency increased, the rate of allele frequency change also increased (Table 4). The most efficient technologies (CRISPR and Perfect) had the fastest rates of change (Fig. 3) and also were the only cases in which observed trends exceeded expected trends (Fig. 4). Differences between methods were much greater than differences between proportions of animals edited. This may be due, in part, to the failure model used in the simulation: when an edit is unsuccessful, the animal’s genotype at the edited locus is not changed. Inefficient technologies will often fail to change heterozygous (*Aa*) genotypes to homozygous dominant (*AA*) genotypes, which reduces the rate at which allele frequencies change. The addition of females did not have a notable effect on rates of allele frequency change. This may be due to the absence from the simulation of advanced reproductive technologies used to propagate elite genotypes (e.g., in vitro fertilization combined with ET for elite cow families to produce flushes of embryos with high PAs).

**Figure 3.**
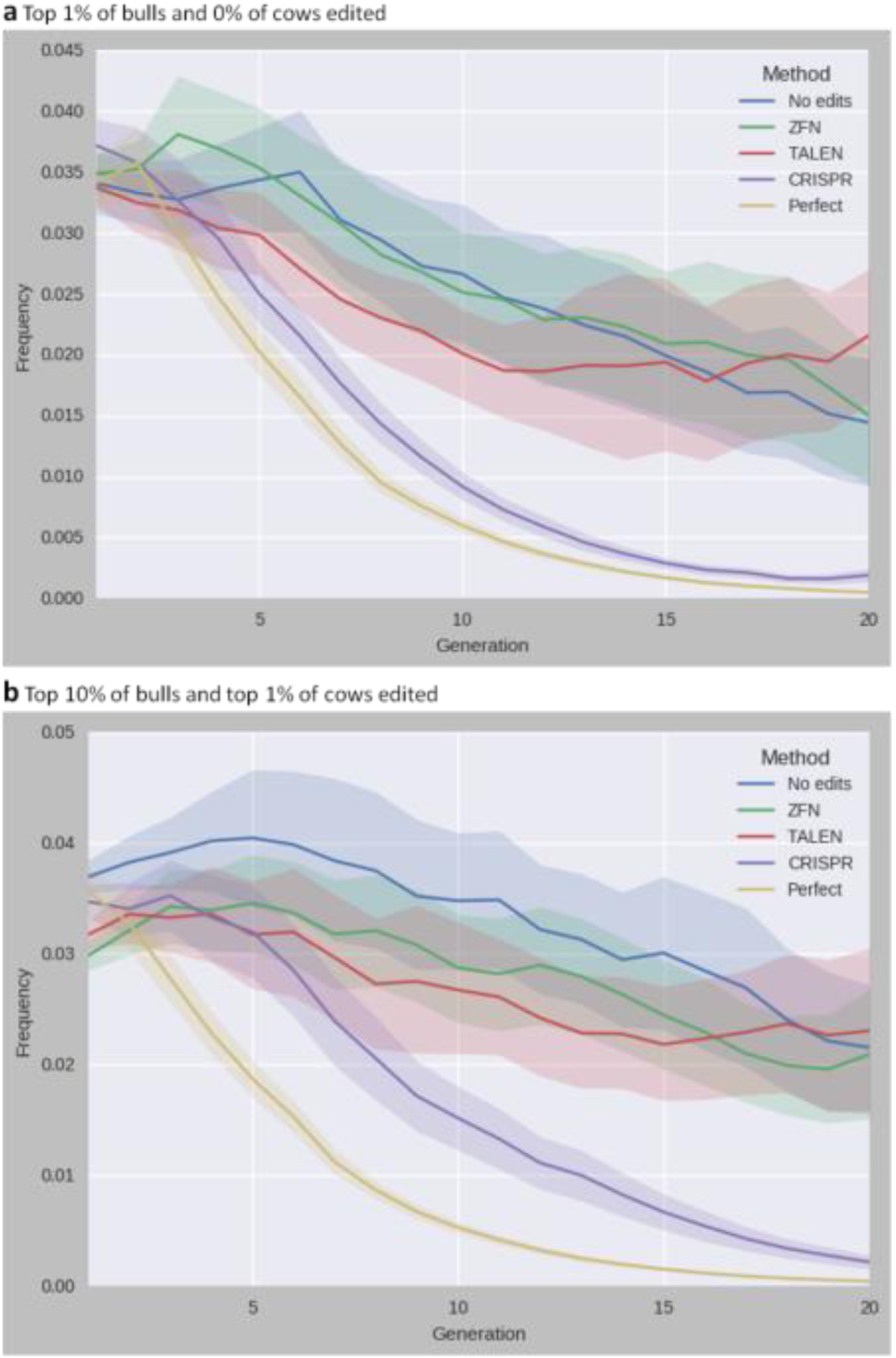
Observed minor allele frequency of the Holstein recessive locus HH3 for five different gene-editing technologies over 20 years. **a** Top 1% of bulls and 0% of cows were edited. b Top 10% of bulls and top 1% of cows were edited. *CRISPR* clustered regularly interspaced short palindromic repeats, *TALEN* transcription activator-like effector nuclease, *Perfect* = hypothetical technology with a perfect success rate, *ZFN* zinc finger nuclease

**Figure 4.**
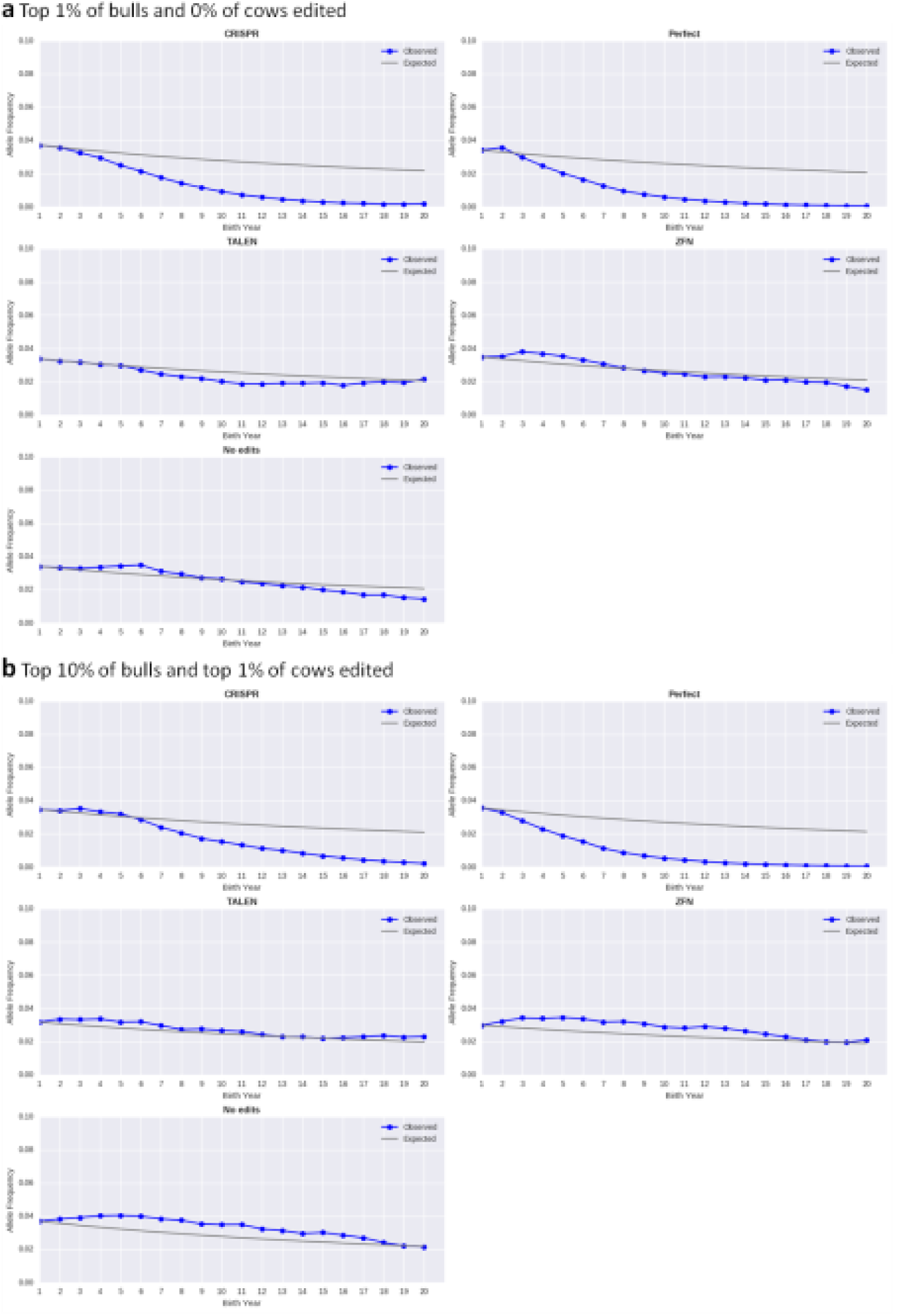
Observed versus expected changes in allele frequencies of the Holstein recessive locus HH3 for five different gene-editing technologies over 20 years. **a** Top 1% of bulls and 0% of cows were edited. b Top 10% of bulls and top 1% of cows were edited. *CRISPR* clustered regularly interspaced short palindromic repeats, *TALEN* transcription activator-like effector nuclease, *Perfect* = hypothetical technology with a perfect success rate, *ZFN* zinc finger nuclease

**Table 4.** Coefficients for regression of observed allele frequency on birth year and standard errors.

| Recessive | Technology | 1% bulls and 0% cows gene edited<sup>a,b</sup> |  |  |  | 10% bulls and 1% cows edited<sup>a,b</sup> |  |  |  |
| --- | --- | --- | --- | --- | --- | --- | --- | --- | --- |
|  |  | b<sub>linear</sub> | SE<sub>linear</sub> | b<sub>quadratic</sub> | SE<sub>quadratic</sub> | b<sub>linear</sub> | SE<sub>linear</sub> | b<sub>quadratic</sub> | SE<sub>quadratic</sub> |
| HH3 | No edits | -0.0007** | 0.000 | **-2.28E-05** | **1.06E-05** | **0.0003** | **0.000** | -5.93E-05** | 8.64E-06 |
|  | ZFN | -0.0014** | 0.000 | **1.059E-05** | **1.3E-05** | **0.0002** | **0.000** | -4.86E-05** | 1.12E-05 |
|  | TALEN | -0.0026** | 0.000 | 8.43E-05** | 8.26E-06 | -0.0012** | 0.000 | **2.677E-05** | **1.07E-05** |
|  | CRISPR | -0.0048** | 0.000 | 0.0001** | 7.67E-06 | -0.0031** | 0.000 | 5.001E-05* | 1.54E-05 |
|  | Perfect | -0.0050** | 0.000 | 0.0002** | 1.13E-05 | -0.0050** | 0.000 | 0.0002** | 8.75E-06 |
| Horned | No edits | -0.0006** | 0.000 | **1.677E-05** | **6.34E-06** | -0.0017** | 0.000 | 0.0001** | 1.89E-05 |
|  | ZFN | -0.0109** | 0.001 | **-0.0002** | **3.66E-05** | -0.0123** | 0.001 | -0.0004** | 6.65E-05 |
|  | TALEN | -0.0303** | 0.003 | **-3.098E-05** | **0.000** | -0.0338** | 0.003 | **0.0001** | **0.000** |
|  | CRISPR | -0.0986** | 0.005 | 0.0022** | 0.000 | -0.1070** | 0.006 | 0.0025** | 0.000 |
|  | Perfect | -0.1428** | 0.006 | 0.0044** | 0.000 | -0.1418** | 0.006 | 0.0044** | 0.000 |

Regression was for five different editing technologies over 20 years in scenarios where either the top 1% of bulls and 0% cows were edited or the top 10% of bulls and top 1% of cows were edited. *CRISPR* clustered regularly interspaced short palindromic repeats, *TALEN* transcription activator-like effector nuclease, *Perfect* hypothetical technology with a perfect success rate, *ZFN* zinc finger nuclease; *b* regression coefficient; SE standard error
<sup>a</sup>*t* test significance for the hypotheses that |b<sub>linear</sub>| = 0 and |b<sub>quadratic</sub>| = 0: \**P* < 0.01, \*\**P* < 0.001; bold terms did not differ from 0.
<sup>b</sup>The software used for analysis displays only three digits, SE < 0.001 were truncated to 0.000, but actual values were not exactly 0.

Rates of inbreeding did not differ between editing methods when only bulls and no cows were edited (Fig. 5a). However, rates of inbreeding did differ when both bulls and cows were edited (Fig. 5b) for CRISPR, TALEN, and ZFN. When the ET process fails, the embryo dies (Fig. 1), which results in the loss of a calf with high-genetic-merit in the next generation because only elite embryos are edited. If the process is inefficient and many embryos die, then animals are used as sires that would otherwise not be selected. The reduction in inbreeding is greatest for the least efficient method (ZFN), followed in order by TALEN and CRISPR. Editing methods with high failure rates result in the selection of parents that would not otherwise have been selected, which reduces within-family matings and inbreeding rates.

**Figure 5.**
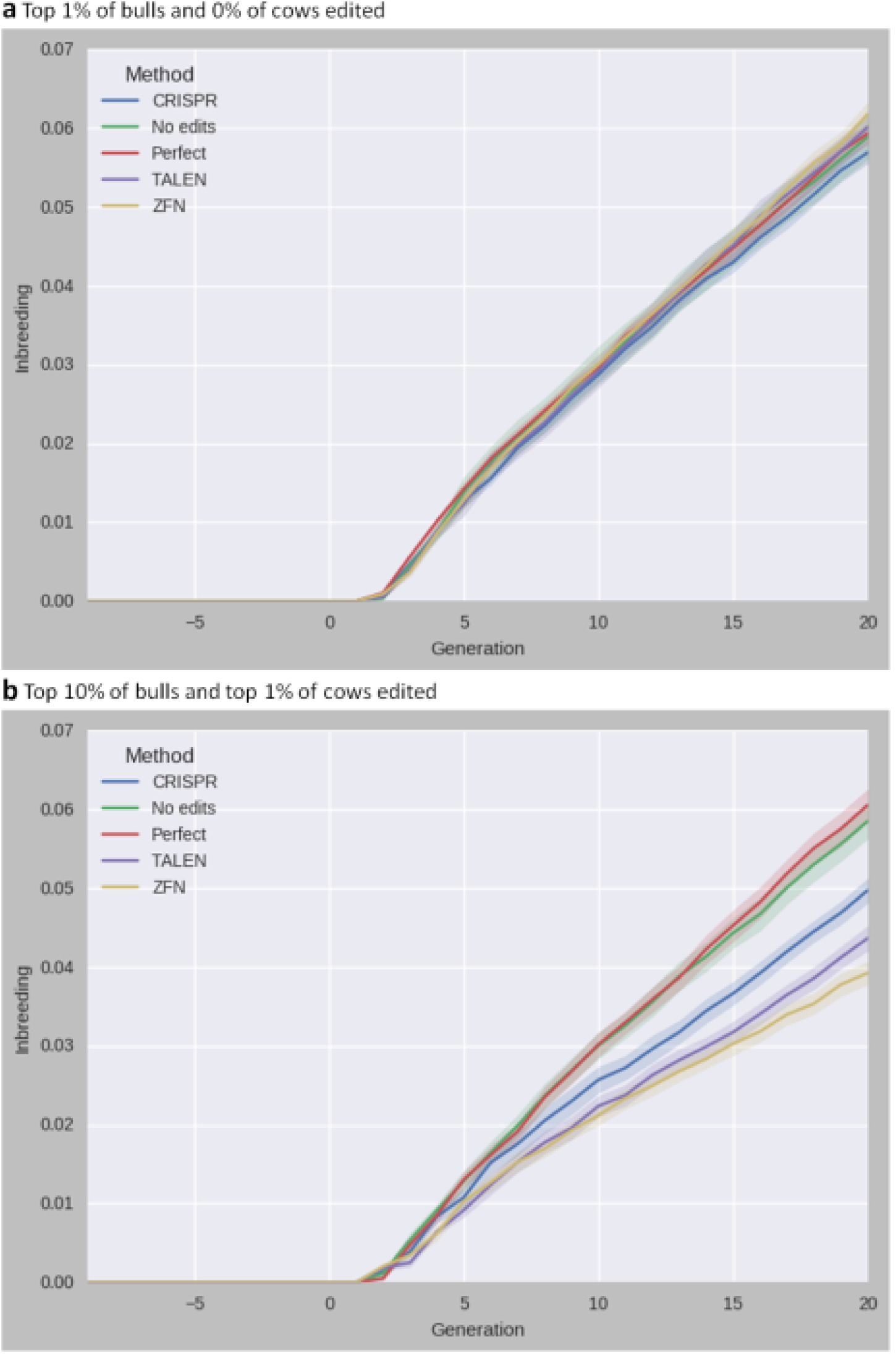
Average inbreeding rate in a simulation of 12 Holstein recessive loci for five different gene-editing technologies over 20 years. **a** Top 1% of bulls and 0% of cows were edited. b Top 10% of bulls and top 1% of cows were edited. *CRISPR* clustered regularly interspaced short palindromic repeats, *TALEN* transcription activator-like effector nuclease, *Perfect* = hypothetical technology with a perfect success rate, *ZFN* zinc finger nuclease

#### Horned

The polled (hornless) state is dominant to the horned state. This discussion is focused on horned, the recessive allele, to mirror the results and discussion for HH3 as well as findings of Cole [5]. Previous studies on breeding strategies for decreasing the frequency of the recessive (horned) allele in dairy cattle (e.g., [5,28,29]) suggested that rates of change would be very slow, and a number of authors have instead proposed selection directly on the polled locus or linked markers (e.g., [30,31,32,33,34]). Long-term progress can be improved slightly by putting more weight on favorable minor alleles in selection programs [35], but progress would be much faster using gene edits for the favorable allele.

In this simulation, a single locus was assumed to control polledness, but in reality the polled locus is more complex than HH3 and has at least two mutations on BTA1 that result in hornless cattle [36,37]. All gene-editing methods resulted in significant rates of allele frequency change (Table 4), with rates of change increasing with the efficiency of the technology (Fig. 6). Regression coefficients were similar regardless of the proportion of bulls and cows edited. In contrast to the results obtained for HH3, observed trends were greater than expected trends for every editing technology (Fig. 7). Differences between methods were much greater than differences between proportions of animals edited. Even the least-efficient editing technology produced large reductions in the frequency of horned cattle, which is a notable improvement over the results of Cole [5], who found no allele frequency change after including the economic value of polledness in the selection criterion. Differences in rates of inbreeding for the horned locus (not shown) were similar to those observed for HH3, again supporting that higher failure rates will result in sampling more diverse pedigrees than would otherwise be the case.

**Figure 6.**
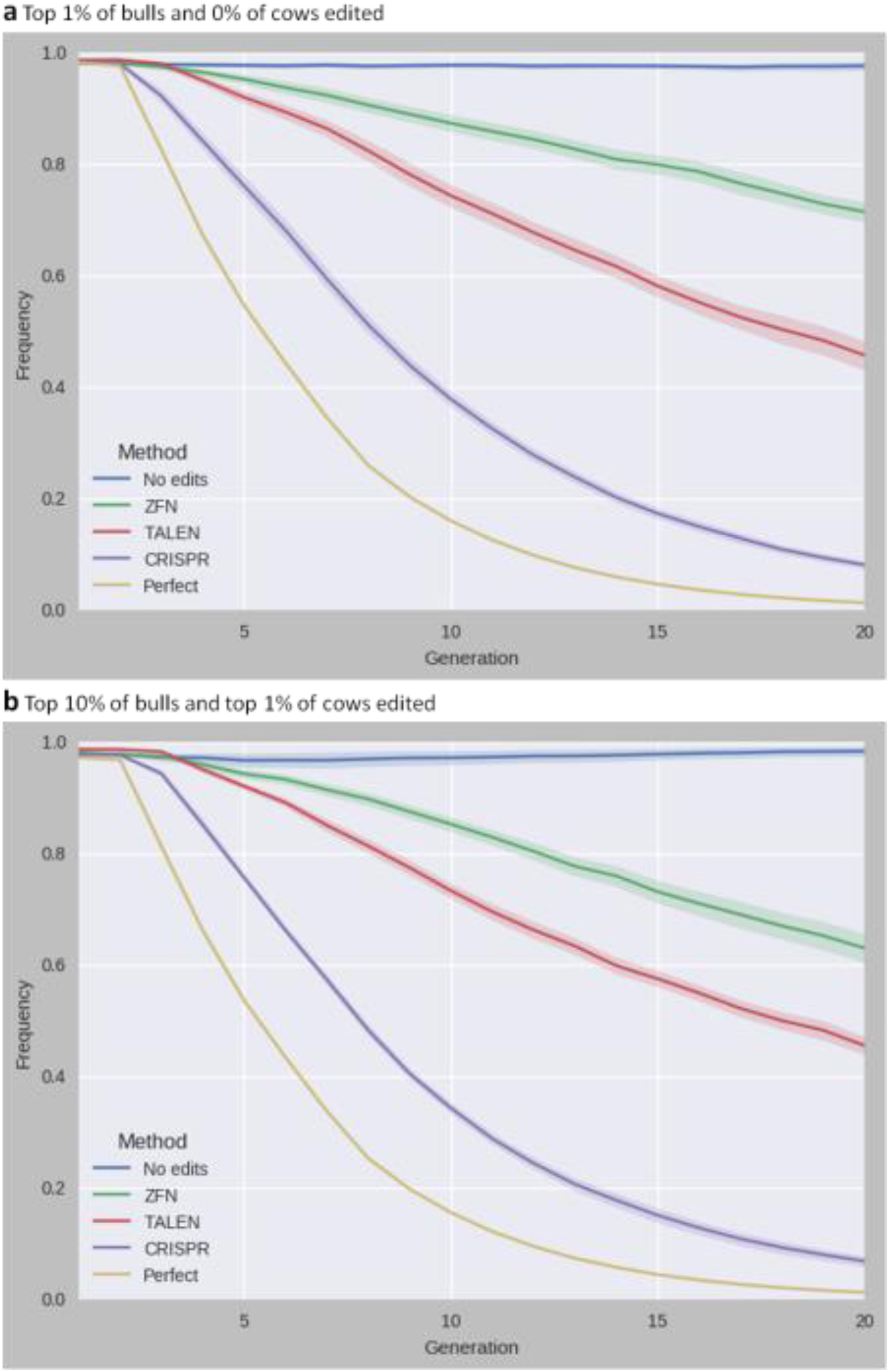
Observed minor allele frequency of the Holstein recessive locus horned for five different gene-editing technologies over 20 years. **a** Top 1% of bulls and 0% of cows were edited. b Top 10% of bulls and top 1% of cows were edited. *CRISPR* clustered regularly interspaced short palindromic repeats, *TALEN* transcription activator-like effector nuclease, *Perfect* = hypothetical technology with a perfect success rate, *ZFN* zinc finger nuclease

**Figure 7.**
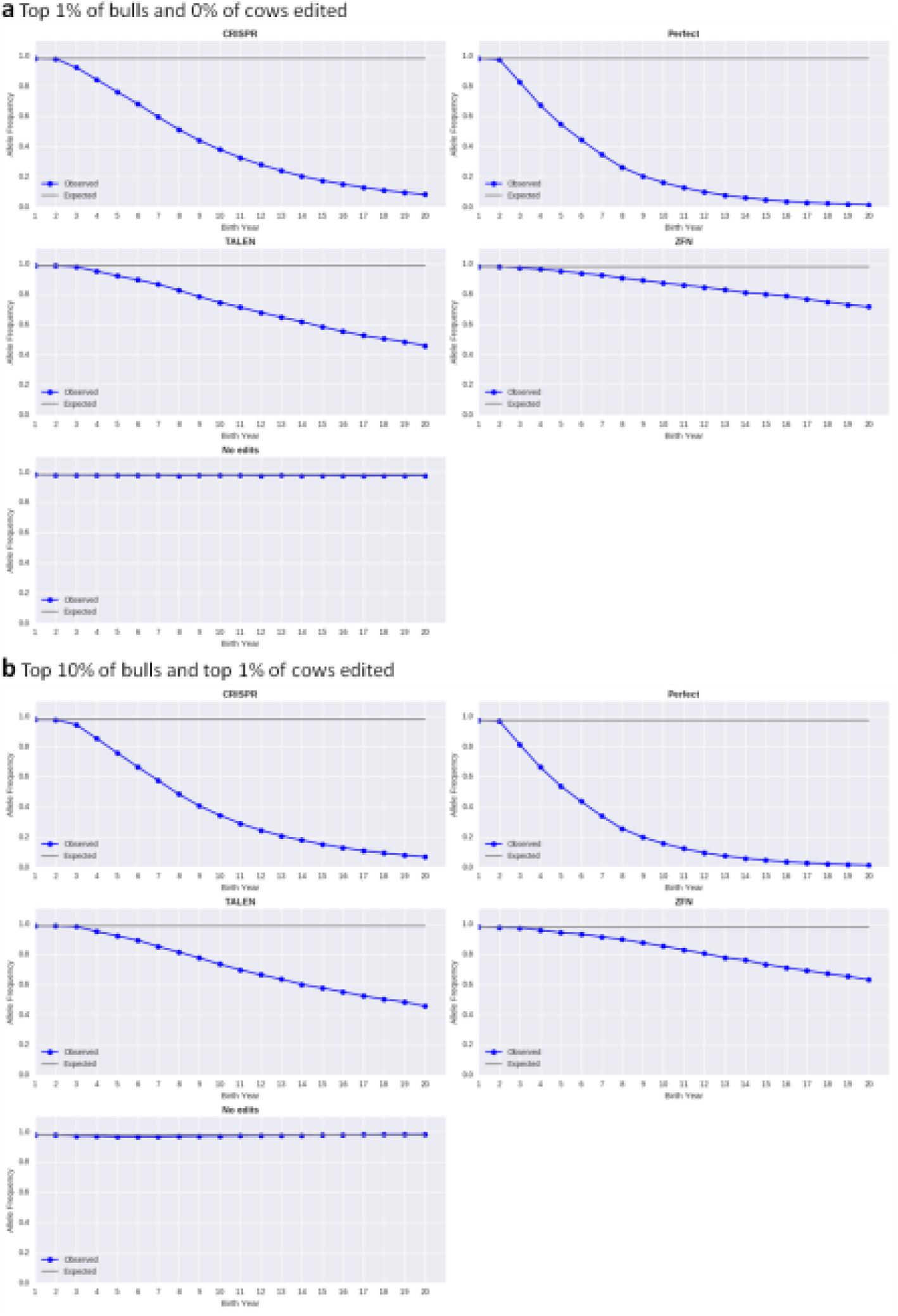
Observed versus expected changes in allele frequencies of the Holstein recessive locus horned for five different gene-editing technologies over 20 years. **a** Top 1% of bulls and 0% of cows were edited. b Top 10% of bulls and top 1% of cows were edited. *CRISPR* clustered regularly interspaced short palindromic repeats, *TALEN* transcription activator-like effector nuclease, *Perfect* = hypothetical technology with a perfect success rate, *ZFN* zinc finger nuclease

#### Cumulative genetic gain

Two sets of *t* tests were conducted to evaluate cumulative genetic gain over the 30 years (10 rounds of burn-in and 20 rounds of selection) of the simulation. First, a *t* test was used to compare each gene editing technology within a scenario against no gene editing to determine if different technologies produce different rates of gain. Then a set of *t* tests was used to compare gene editing technologies across scenarios to determine if the proportion of animals edited had an effect on cumulative gain. Results differed slightly between the horned-only and 12-recessive scenarios; however, the pattern of responses was the same, and only results for the 12-recessive scenario are discussed.

The pattern for cumulative genetic gain was similar to that for rates of inbreeding. The Perfect technology did not differ from no gene editing for either 1% bulls and no cows edited or 10% of bulls and 1% of cows edited. However, CRISPR, TALEN, and ZFN all showed significantly lower cumulative genetic gains (*P* < 0.01), with larger differences for less efficient technologies. Similarly, the scenarios with higher rates of editing also had no differences for no gene editing and the Perfect technology as well as significantly lower rates of gain for CRISPR, TALEN, and ZFN (*P* < 0.01). As previously discussed, when many embryos die during ET, fewer elite animals are available to become parents in the next generation. This resulted in lower rates of genetic gain that were proportional to the ET failure rate over the course of the simulation.

#### Scenario comparison

For both editing scenarios (1% of bulls and 0% of cows edited, and 10% of bulls and 1% of cows edited), the use of gene editing resulted in faster allele frequency changes; more efficient technologies produced faster rates of change. A comparison of results for the horned locus from the horned-only and 12-recessive scenarios (not shown) indicates that the number of loci edited in a scenario had no effect on the rates of allele frequency change. This is expected because gene edits are modelled as independent events, and few animals are carriers of more than one recessive.

### Embryonic losses

The proportion of embryos that died in each birth year because they were homozygous for lethal conditions (Fig 8) also decreased rapidly when only 1% of bulls and no cows were edited using CRISPR and TALEN. When the editing procedure is highly efficient, fewer affected embryos are produced, even when the number of edited parents is few. The Perfect technology produced rapid decreases in both scenarios as expected.

**Figure 8.**
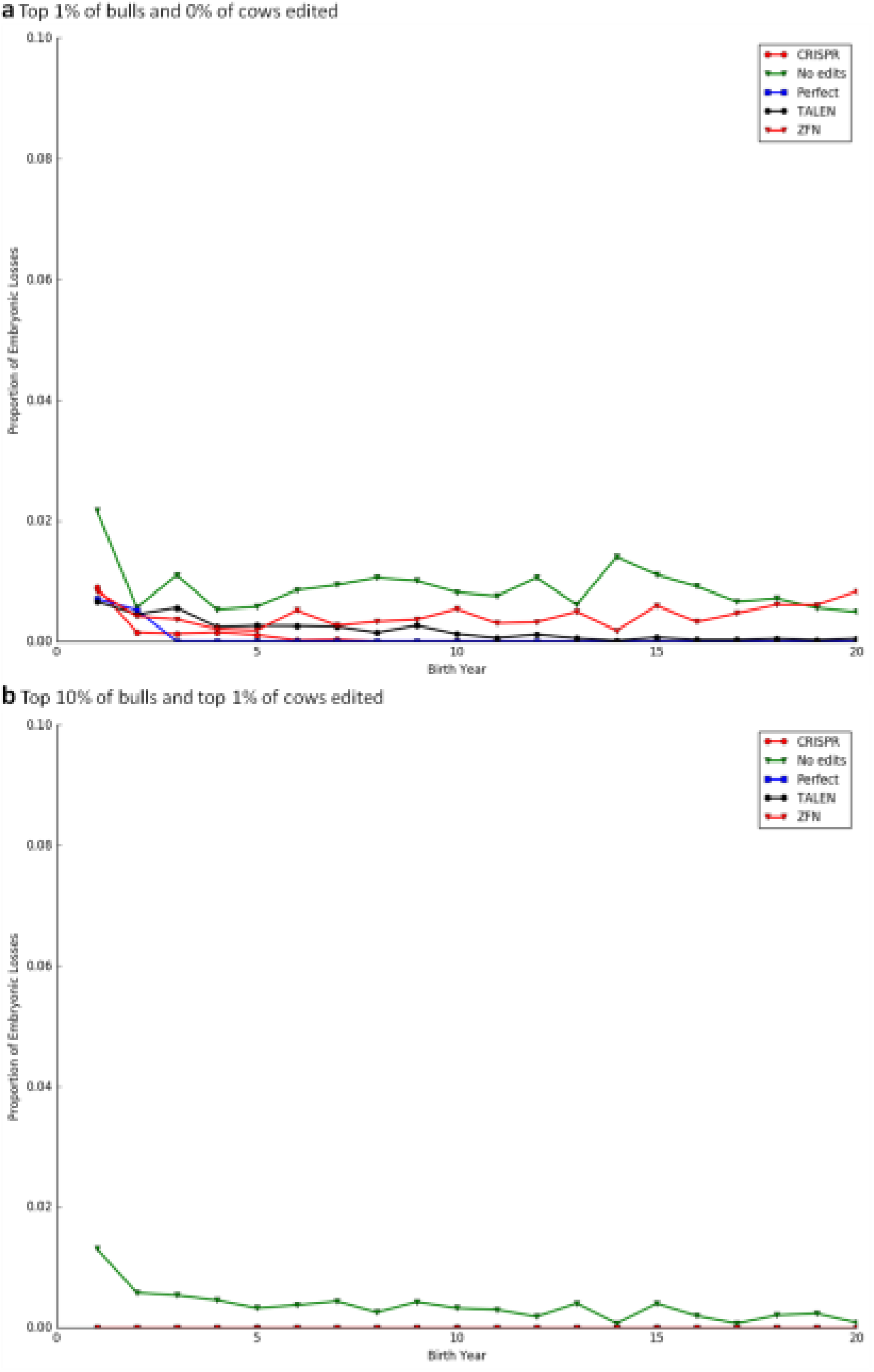
Proportion of embryos that died each year because of the effects of recessive genotypes for five different gene-editing technologies over 20 years. **a** Top 1% of bulls and 0% of cows were edited. b Top 10% of bulls and top 1% of cows were edited. *CRISPR* clustered regularly interspaced short palindromic repeats, *TALEN* transcription activator-like effector nuclease, *Perfect* = hypothetical technology with a perfect success rate, *ZFN* zinc finger nuclease

These results contrast with those of Cole [5], who reported that the number of embryos that died from recessive disorders was higher for scenarios with constrained inbreeding and penalized carrier parents. Cole concluded that the goal of eliminating recessive alleles from the population (fixing haplotypes in a homozygous state) conflicted with the goal of minimizing inbreeding (avoidance of increases in homozygosity). However, if loci can be edited, then favourable alleles can be introduced without affecting overall inbreeding in the population because the number of known recessives is low compared to the size of the genome. These results also may reflect the parameters used in the simulation. With a cow population of 100,000 and 5000 matings permitted per year per bull, a cohort of only 20 bulls could breed every cow in the population. If the top 10% of bulls (*n* = 50) were edited, all cows could have been bred to gene-edited bulls and no affected embryos produced.

### Avoidance of carrier bulls

The proportion of bulls edited had only small effects on allele frequency changes, and the proportion of cows edited had essentially no effect on allele frequency changes, possibly because of the small proportion of cows edited. However, assuming higher editing rates is unrealistic given current limitations of the technology. In the future, artificial-insemination firms may simply refuse to purchase embryos or calves that are carriers of known genetic defects and forgo very little (or no) genetic gain to eliminate recessives from the population rapidly. Cole et al. [3] showed that genetic merit for NM$ of Ayrshire and Jersey bulls that carry at least one known recessive disorder does not differ from those free of known recessives and that Brown Swiss (*P* = 0.087) and Holstein (*P* < 0.001) carriers have lower average predicted transmitting abilities than non-carriers. The proportion of genotyped Holstein bulls and cows born since 2000 that are carriers of at least one known recessive was fairly constant between 2000 and 2010 but began to decrease more quickly in 2011(Fig. 9), which is when haplotype tests were introduced for several new genetic disorders [1]. Collectively, these results suggest that artificial-insemination firms already avoid carrier bulls when purchasing embryos and young bulls. This approach is probably the fastest and least expensive for eliminating harmful genetic defects from the population, but it will not increase the frequency of desirable attributes (e.g., polledness).

**Figure 9.**
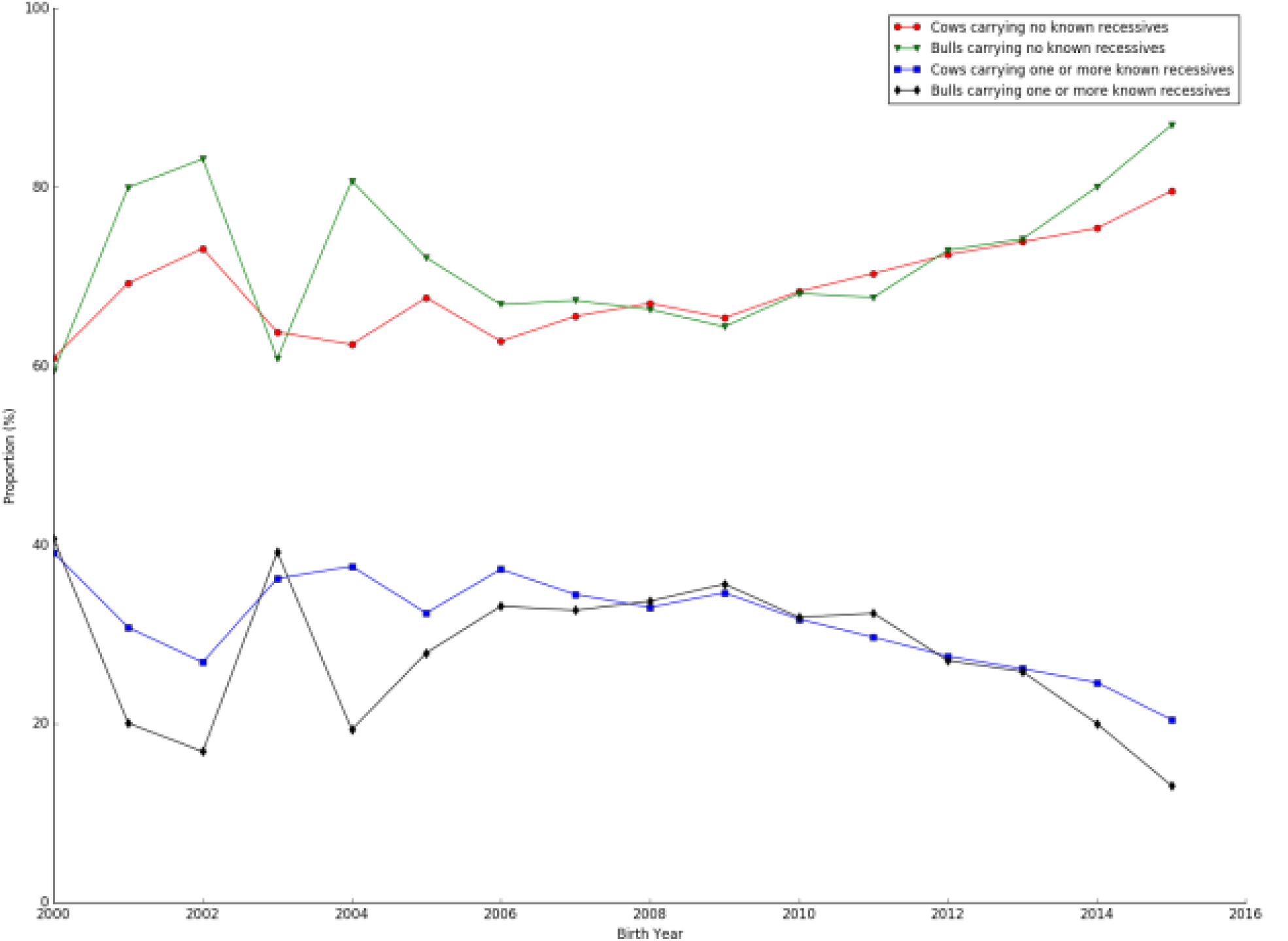
Proportions of genotyped Holsteins that have known recessives by animal sex. Bulls and cows were born from 2000 through 2015. Carrier status was either not a carrier of a known recessive or a carrier of one known recessive or more.

### Regulatory considerations

Although some gene-edited products recently have reached the U.S. marketplace [38,39], regulatory uncertainty remains a concern [40]. This was underscored by much of the discussion at the 2016 Large Animal Genetic Engineering Summit [41], which focused largely on the use of gene editing and other tools to produce large animal models of human disease (e.g., [42]) rather than food animals because of the concerns surrounding consumer acceptance of gene-edited animals in the food chain. Policymakers and regulators are being encouraged to exercise oversight based on the product rather than the tool used to generate that product [43], but if (or when) meat and milk from gene-edited animals will be offered for sale is not clear at present.

### Acceptance of gene editing

Consumers and regulators may be more willing to support the use of gene editing for improving animal welfare rather than simply for increasing productivity. For example, the process of dehorning is traumatic to calves, unpleasant for farmers, and distasteful to consumers (e.g., [44]). Previous studies [5,29] have shown that increasing the frequency of polled animals in the Holstein population is difficult because the frequency of the dominant allele is very low (0.0061). Carlson et al. [11] have successfully produced polled clones of horned animals using gene editing with no detectable off-target effects, which shows that the technology could be used to propagate desirable polled genotypes rapidly. Gene editing also has been used to produce animals with increased resistance to disease [45], including porcine reproductive and respiratory syndrome [10,46] and bovine tuberculosis [47]. Other candidates for gene editing include casein variants that may have beneficial effects on human health [48], the *slick* locus that is involved in adaptation to hot environments thermotolerance [49], and the *DGAT1* gene which has favourable effects on milk composition [50].

Many challenges are associated with genetically modified organisms, some technical and others related to consumer attitudes towards the technology [51,52]. Although the technology has improved dramatically in recent years, the general public remains concerned about genetically modifying food crops and livestock. A recent meta-analysis of the literature on consumer preferences suggests that U.S. respondents have a more favourable view of biotechnologically modified food products than those from Europe, but that most consumers are concerned about genetically modified animals [53]. Consumers that are generally opposed to the marketing of genetically modified organisms may moderate those opinions in the presence of another benefit (e.g., increased levels of omega-3 fatty acids in farmed salmon) [54]. Changing consumer attitudes towards technologies may be possible, but the arguments need to focus on the benefits rather than the technology [55]. Consumers may be more accepting of gene editing in food animals if the technology focus is on animal health and welfare rather than productivity.

## Conclusions

The efficiency of gene editing technologies has a greater effect on allele frequency change than the proportion of animals in the population edited. Gene editing is a useful tool for increasing the frequency of desirable characteristics that are at low frequency in current populations (e.g., polledness). Removing carriers of harmful recessives from the population may be more effective than correcting them with gene editing. The use of gene editing to increase the frequency of alleles that confer resistance to disease may be more acceptable to consumers than using the technology to increase genetic merit for quantitative traits. Applications of gene editing in livestock should focus on loci with large, beneficial impacts on animal health rather than on recessive defects with low allele frequencies in the population, because the latter can better be managed through mating programs.

## Declarations

### Ethics approval and consent to participate

This study involved no animal experimentation and did not require any authorization from local ethics committee.

### Consent for publication

Not applicable.

### Availability of data and material

The source code for the simulation and the Jupyter [**56**] notebooks used to analyze the data are available on GitHub [**14**] under a Creative Commons CC0 1.0 Universal (public domain) license.

### Authors’ contributions

Not applicable.

### Acknowledgements

Suzanne Hubbard and Daniel Null (Animal Genomics and Improvement Laboratory, Agricultural Research Service, USDA, Beltsville, MD) assisted with figures and technical editing, and Kristen Parker Gaddis (Council on Dairy Cattle Breeding, Bowie, MD) and Paul VanRaden (Animal Genomics and Improvement Laboratory, Agricultural Research Service, USDA, Beltsville, MD) provided several insightful comments about the manuscript.

